# Mutational signatures in cancer genomes alter protein sequence motifs in cellular signaling networks

**DOI:** 10.64898/2026.02.03.703397

**Authors:** Jigyansa Mishra, Masroor Bayati, Zoe P. Klein, Kevin C. L. Cheng, Omar Wagih, Nina Adler, Maria Farina-Morillas, Alexander T. Bahcheli, Jüri Reimand

## Abstract

Somatic mutations in cancer genomes arise from distinct, context-specific mutational processes, yet their functional consequences at the protein and network level remain incompletely understood. Here, we show that mutational processes of single-nucleotide variants (SNVs) can systematically rewire signaling networks by inducing amino acid substitutions in short linear motifs (SLiMs) that mediate interactions with kinases and other signaling proteins. By analysing 11,000 cancer genomes and 144 classes of SLiMs, we identify motif-rewiring SNVs (rwSNVs) that create or disrupt SLiMs or remove phosphorylated residues. Mutational processes of methylcytosine deamination, APOBEC activity, and ultraviolet light exposure emerge as major contributors to motif rewiring. rwSNVs are enriched in cancer driver genes and pathways, linking mutation etiology to functional consequences. rwSNVs at the BRAF V600E hotspot associated with UV-related mutagenesis are predicted to generate a phosphorylation motif recognized by PLK1 kinase. Together, these findings reveal how mutational processes shape oncogenic signaling and tumor heterogeneity.

## Introduction

Mutational processes shape cancer evolution by generating genomic alterations in a tissue- and context-specific manner ^1,2^. Single-nucleotide variants (SNVs), the most common class of mutations in cancer genomes, arise from diverse endogenous and exogenous mutational processes that preferentially generate substitutions in specific trinucleotide contexts, giving rise to characteristic signatures of single-base substitutions (SBS) ^3^. SBS signatures have been linked to clock-like mutagenesis and aging ^4^, carcinogenic exposures such as tobacco smoke ^5^ and UV light ^6^, defects in DNA repair ^7^, APOBEC enzyme activity ^8^, and cancer therapies ^9^. While most alterations in cancer genomes are considered functionally neutral passengers, a few alterations confer selective advantages to cells and act as cancer drivers ^10,11^.

The genomic footprints of mutational processes suggest that their sequence biases may have systematic consequences for the proteome. Because mutational processes tend to alter DNA in specific trinucleotide contexts, these biases can also alter the spectrum of amino acid substitutions introduced in proteins through the triplet codon structure of the genetic code. Previously, we showed that nonsense SNVs are strongly associated with mutational signatures of APOBEC activity and tobacco smoking, and corresponding nonsense SNVs cause inactivation of tumor suppressor genes, implicating these mutational processes in loss-of-function mutagenesis ^12^. It remains unclear whether comparable biases of mutational processes exist for missense SNVs that could systematically alter proteins and cellular processes in cancer.

Cellular signalling networks regulate biological processes through dynamic, modular protein-protein interactions, many of which are mediated by short linear motifs (SLiMs) ^13^. SLiMs are compact sequence elements that recruit signalling proteins, for example motifs bound by kinases to catalyse phosphorylation events, and motifs bound by SH2, SH3, or 14-3-3 domains that mediate phosphorylation-dependent protein-protein interactions central to signal transduction ^14^. Many SLiM-mediated interactions occur within intrinsically disordered regions (IDRs) of proteins where they enable selective yet transient binding events that organize and propagate cellular signals ^15^. Large-scale proteomics studies have identified hundreds of thousands of phosphorylation sites, a substantial fraction of which reside within known SLiMs ^16,17^. Phosphorylation signaling is central to hallmark cancer pathways, and kinases constitute one of the largest and most therapeutically targeted protein families in oncology ^18^.

As SLiMs and phosphorylation sites are short and defined by a small number of critical residues, they are particularly sensitive to protein-coding mutations in cancer genomes. Missense SNVs can therefore create new SLiMs or disrupt existing ones, potentially altering signaling interactions ^19,20^. A well-established example is the β-catenin oncogene *CTNNB1*, in which recurrent mutations accumulate within N-terminal phosphorylation motifs, impairing protein degradation and causing aberrant activation of Wnt signaling in cancer ^21,22^. More broadly, phosphorylation sites are under purifying selection in the human population and are enriched for inherited disease variants as well as somatic driver mutations in cancer genomes ^23,24^. Together, these observations highlight the susceptibility of SLiM-mediated signaling interactions to genetic variation and underscore their potential role in pathogenic rewiring of cellular signaling networks.

Here, we test the hypothesis that mutational processes active in cancer genomes systematically rewire cellular signaling networks by preferentially disrupting or creating short linear motifs. Because mutational signatures are enriched for mutations within specific trinucleotide contexts, and therefore for specific amino acid substitutions, they are expected to differentially impact distinct classes of sequence motifs. To examine this, we analyzed missense SNVs across primary and metastatic tumors, predicted their motif-rewiring effects, and assessed their associations with mutational signatures. We find that distinct mutational processes selectively and pervasively alter SLiMs in cancer genomes, establishing a mechanistic link between somatic mutagenesis, perturbation of signaling networks, and tumor heterogeneity.

## Results

### Mutational processes alter phosphosites and SLiMs via rwSNVs

To examine how mutational processes impact cellular signaling networks, we integrated somatic mutations from cancer genomes with SLiMs in protein sequences (**Figure 1a**). We analyzed 124 classes of kinase-bound SLiMs mapped to 149,275 experimentally validated phosphorylation sites ^17,20^, together with 20 additional signaling-related SLiMs from the ELM database ^25^ (including SH2, SH3, and 14-3-3 binding motifs) mapped to IDRs of proteins. In total, we analysed 1.24 million missense SNVs from 11,962 cancer genomes across 19 cancer types, comprising 4,992 primary tumors from The Cancer Genome Atlas (TCGA) ^26^ and 6,970 metastatic tumors from the Hartwig Medical Foundation (HMF) ^27^. Each SNV was assigned to an SBS mutational signature using SigProfilerAssignment ^28^.

**Figure 1.**
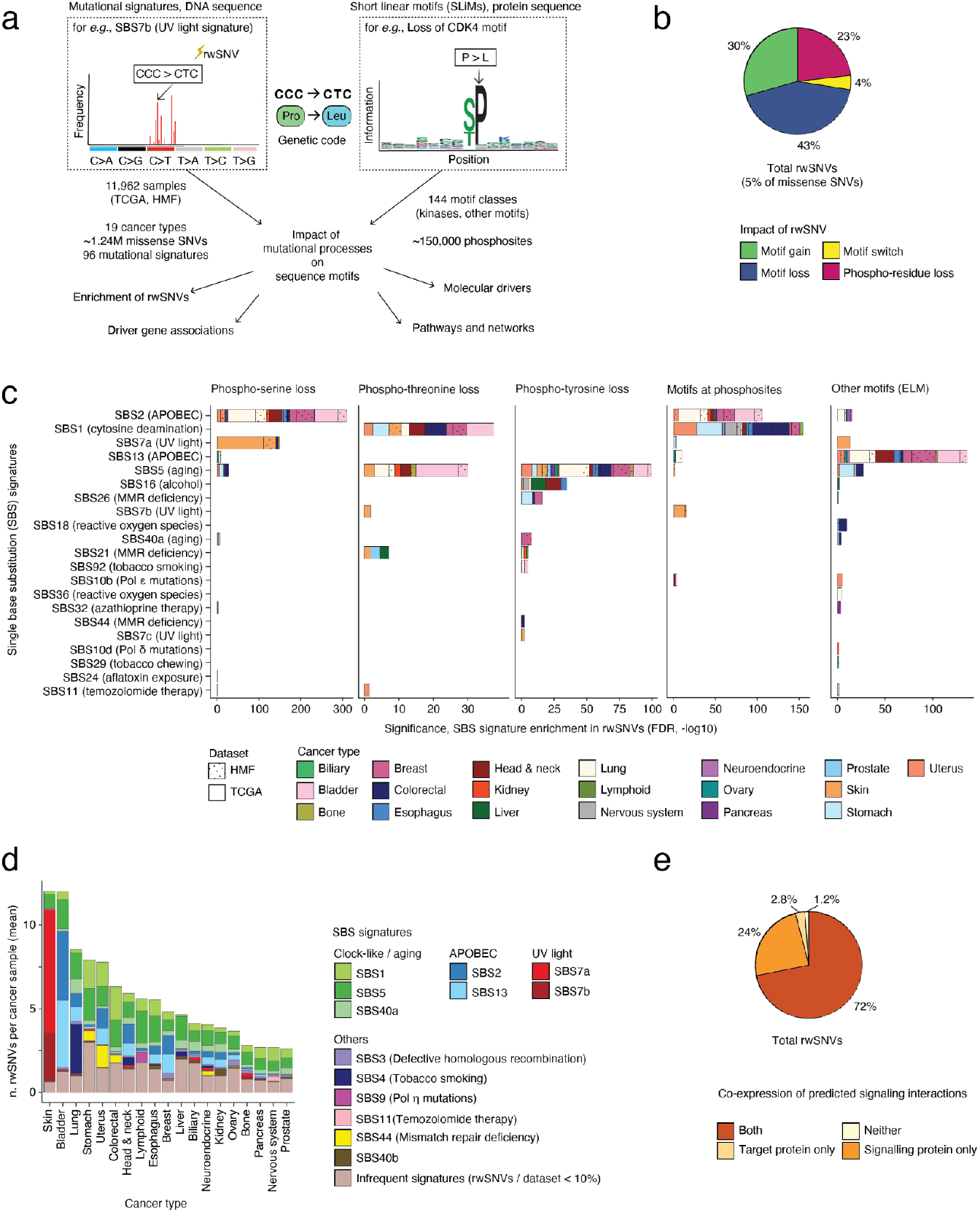
Mutational processes in cancer genomes alter protein SLiMs. **(a)** Overview of the analysis integrating missense SNVs from cancer genomics datasets with SLiMs and mutational processes. **(b)** Distribution of motif-rewiring SNVs (rwSNVs) by predicted SLiM impact. **(c)** Enrichments of SBS mutational signatures in rwSNVs for individual cancer types (shown as colors) and the two genomics cohorts TCGA and HMF (hypergeometric tests, FDR < 0.05). rwSNV enrichments are grouped by functional impact on phosphorylation sites or SLiMs (left to right). **(d)** Mean rwSNV burden in samples of different cancer types, grouped by SBS signatures inducing the rwSNVs (shown in colors). **(e)** Co-expression analysis of rwSNV-affected proteins and the SLiM-associated signaling proteins based on matching transcriptomics data in TCGA. The pie chart shows the proportions of signaling interactions between signaling proteins and target proteins that are supported by gene expression in the cancer samples in which the relevant SLiMs and rwSNVs were identified.

We defined a subset of missense SNVs as rwSNVs, comprising variants that either replace experimentally validated phosphoresidues or cause gains, losses, or switches of SLiMs as predicted by the machine learning method MIMP ^20^. Across all cancer types, we identified 62,850 rwSNVs, corresponding to approximately 5% of all missense SNVs (posterior p > 0.8 from MIMP; **Figure 1b, Table S1**). Among these, 23% of rwSNVs directly replaced phosphorylated residues (Ser, Thr, or Tyr) with non-phosphorylatable amino acids. The majority of rwSNVs altered critical residues within SLiMs, resulting in motif losses (43%), motif gains (30%), or motif switches involving concurrent gains and losses of different SLiMs (4%). Direct substitution of phosphoresidues is expected to abolish post-translational modification at the site, whereas SLiM-altering rwSNVs are predicted to reconfigure local signaling interactions.

We next asked which mutational processes were most strongly associated with rwSNVs across cancer types. In total, we identified 190 significant enrichments stratified by rwSNV functional impact (one-tailed hypergeometric tests, FDR < 0.05; **Figure 1c, Table S2**). APOBEC-associated signatures (SBS2 and SBS13) and clock-like signatures (SBS1 and SBS5) showed the most consistent enrichment in rwSNVs across multiple cancer types, particularly in breast, bladder, lung, and colorectal cancers. In addition, tissue-specific mutational processes were associated with rwSNVs, including UV light mutagenesis in melanoma (SBS7a), alcohol-associated mutagenesis in liver and head and neck cancers (SBS16), and reactive oxygen species-associated mutagenesis in colorectal cancer (SBS18).

Distinct mutational signatures were associated with different classes of rwSNV functional impact. The APOBEC signature SBS2 was preferentially enriched in phosphoserine losses (*e*.*g*., primary breast cancer, FDR = 1.75 x 10^-17^, OR = 2.8) and alterations of phosphosite SLiMs (*e*.*g*., primary lung cancer, FDR = 4.7 x 10^-26^, OR = 2.0), whereas SBS13 more frequently altered SLiMs from the ELM database (*e*.*g*., metastatic bladder cancer, FDR = 1.9 x 10^-8^, OR = 1.7). In contrast, clock-like signatures SBS1 and SBS5 were enriched in losses of phosphothreonine and phosphotyrosine residues, respectively.

We next quantified motif-rewiring mutations across cancer genomes by analysing rwSNV burden per cancer sample (**Figure 1d**). On average, we found 5.3 rwSNVs per cancer genome, with substantial variation across cancer types. Melanoma and bladder cancer samples exhibited the highest rwSNV burden with 12 rwSNVs per sample on average, consistent with their elevated overall mutation rates, whereas the fewest rwSNVs were found pancreatic, prostate, and central nervous system cancers (2.6 rwSNVs on average). Tissue-specific contributions of mutational processes to rwSNV burden were also evident. In melanoma, 80% of rwSNVs per sample were attributable to UV-associated mutagenesis, whereas APOBEC-driven processes contributed to 8.1 rwSNVs per sample in bladder cancer and 1-2 rwSNVs per sample in breast, lung, head and neck, and uterine cancers. These results illustrate the extent to which diverse mutational processes contribute to motif-rewiring alterations across cancer types.

To assess the functional relevance of rwSNVs, we evaluated whether rwSNV-affected proteins and their predicted signaling interaction partners were co-expressed at the mRNA level in corresponding cancer samples. Using matched transcriptomic data from TCGA, we found that 72% of rwSNVs were supported by co-expression of both the target protein and the signaling proteins predicted to interact with the affected SLiMs (**Figure 1e, Figure S1a-c**). In 24% of rwSNVs, the interacting signaling proteins were expressed whereas the target proteins were not detectably expressed. Other determinants of signaling interactions, such as protein abundance and subcellular localisation, were not assessed here. Nevertheless, the widespread co-expression of rwSNV-associated protein pairs supports the functional relevance of these predicted motif-rewiring events.

### Mutational processes of APOBEC, cytosine deamination and UV exposure alter critical positions in SLiMs

Because SLiMs depend on a small number of critical residues, their creation or disruption can result from a limited set of amino acid substitutions. The trinucleotide preferences of distinct mutational processes can therefore shape which signalling network modules are most frequently affected. To examine this, we asked which classes of SLiMs were preferentially targeted by rwSNVs from individual mutational signatures and visualized their associations as a network, collapsing overlapping rwSNV sets into non-redundant subnetworks.

Across the cancer types analyzed, we identified 251 significant enrichments of mutational signatures causing rwSNVs in specific SLiMs (FDR < 0.05, one-tailed hypergeometric tests, **Figure 2a, Table S3**). Mutational signatures exhibited distinct functional preferences, with some predominantly causing phosphoresidue losses and others inducing gains or losses of SLiMs (**Figure 2b**). We therefore examined the major rwSNV-generating mutational processes in more detail.

**Figure 2.**
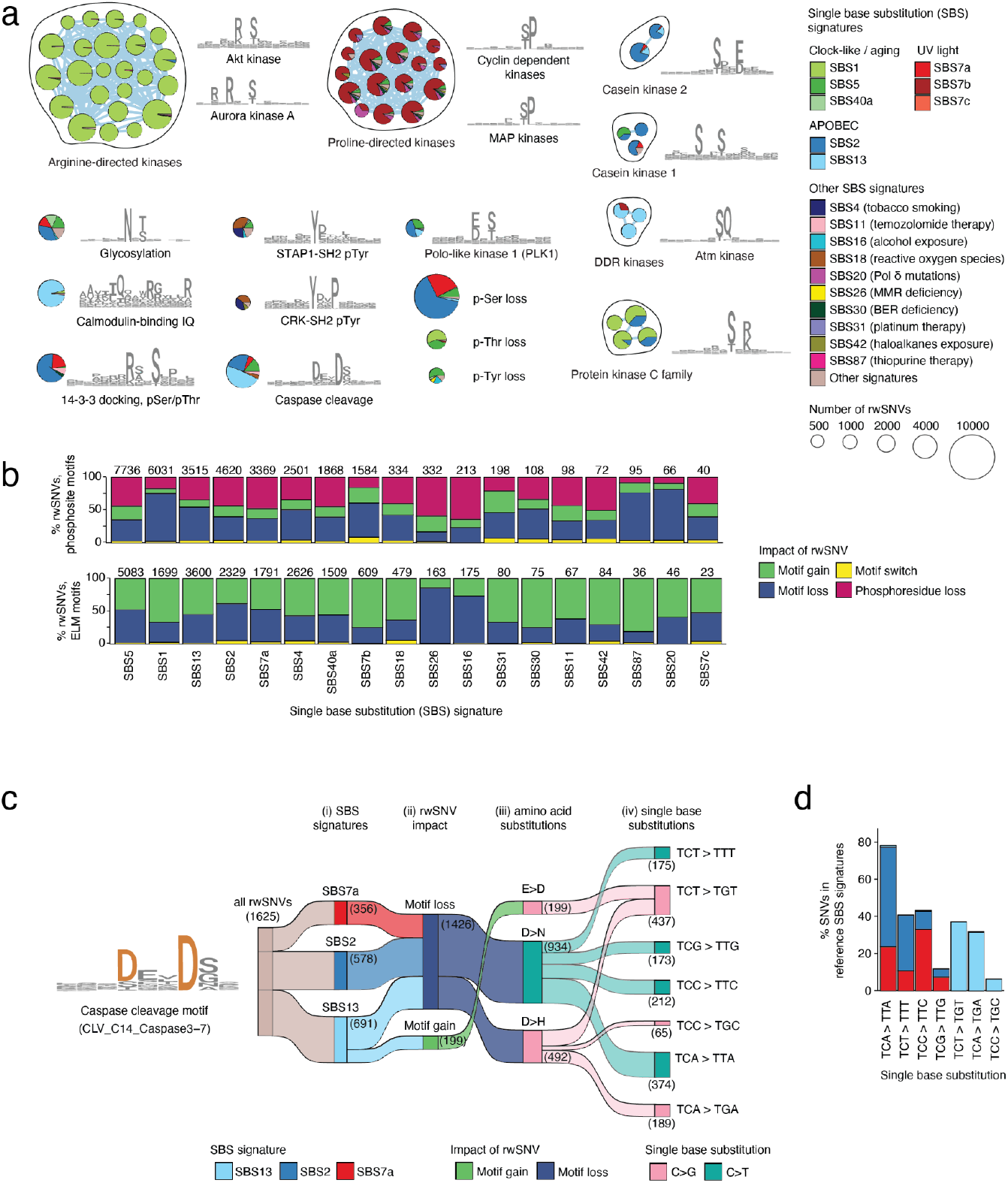
Mutational processes drive distinct genetic alterations of SLiMs. **(a)** Associations of mutational signatures and SLiM classes altered by rwSNVs in multiple cancer types. Significant associations (hypergeometric tests, FDR < 0.05) are visualized as an enrichment map where each node represents a SLiM that is enriched in rwSNVs of a given mutational signature. Phosphoresidue losses are indicated as additional nodes. Nodes are connected by edges if the corresponding SLiMs share some rwSNVs (*i*.*e*., multiple SLiMs are affected by the same subset of rwSNVs). Node size corresponds to rwSNV counts (log-scale) and piecharts reflect significance of mutational signatures contributing to rwSNVs. For each subnetwork, representative SLiMs are visualized as position weight matrix logos. SBS signatures with at least two significant SLiM associations are included and other signatures are collapsed (shown in beige). Associations involving at least 500 rwSNVs are shown. **(b)** Functional impacts of mutational processes on SLiMs. The bar plot shows the rwSNVs from each mutational signature grouped by motif gain, loss, or switch, or loss of phosphoresidue. Total rwSNV counts from all cancer samples are shown above the bars. **(c)** Motif-altering impact of rwSNVs in the caspase cleavage motif (left) is explained by the trinucleotide content of SBS signatures and genetic code. The Sankey diagram (right) shows how three major SBS signatures including APOBEC (SBS2, SBS13) and UV light (SBS7a) interact with amino acid substitutions altering the caspase cleavage motif. Steps of SBS signatures rewiring SLiMs are shown: (i) SBS signature contributions; (ii) rwSNVs causing SLiM gains or losses; (iii) corresponding amino acid substitutions causing SLiM alterations; (iv) corresponding nucleotide substitutions in trinucleotide contexts. Infrequent trinucleotides contributing to rwSNVs at this SLiM are not shown (<1%). **(d)** Frequencies of the main rwSNV-encoding trinucleotides of the three SBS signatures from panel **c**, based on the canonical SBS signature definitions from the COSMIC database.

The clock-like signature SBS1 representing methylcytosine deamination ^7^ was strongly associated with rwSNVs altering basophilic phosphorylation motifs across multiple cancer types (**Figure 2a, Figure S2a**). These motifs contain arginine residues (*e*.*g*., X-R-X-[pS/pT], R-X-X-[pS/pT]) and are bound by AGC family kinases (*e*.*g*., PKA, AKT, PKC), calcium/calmodulin-dependent kinases (CaMKs) and checkpoint kinases (CHK1, CHK2), which are involved in key pathways controlling cell survival and proliferation, metabolism, and stress responses ^16,29^.

APOBEC-associated signatures SBS2 and SBS13 affected diverse SLiMs across cancer types with distinct rwSNV-based impacts (**Figure 2a**). SBS2 was enriched in losses of casein kinase motifs (CK1: S-X-X-[pS/pT]; CK2: [pS/pT]-X-[E/D]) (**Figure S2d-e**), whereas SBS13 more frequently altered caspase-cleavage motifs (D-X-X-D), calmodulin-binding motifs, and kinase-bound SLiMs involved in DNA damage response pathways ^30^ (**Figure S2b**).

To illustrate the genetic basis of these associations, we examined APOBEC-driven rwSNVs affecting the caspase-cleavage motif D-X-X-D (**Figure 2c**). SBS2 predominantly induced T[C>T]N nucleotide substitutions that led to D>N amino acid substitutions, causing disruption of caspase cleavage motifs across multiple cancer types. A similar effect was also observed for the UV signature SBS7a in melanoma. In contrast, SBS13 induced T[C>G]N nucleotide substitutions, resulting in E>D amino acid substitutions that created new caspase motifs, or D>H amino acid substitutions that disrupted existing motifs. These patterns reflect the dominant trinucleotide contexts of SBS2 and SBS13 in the COSMIC database ^31^ (**Figure 2d**), linking mutational processes to motif-level functional consequences.

UV-associated mutational signatures further illustrated process-specific motif rewiring. In melanoma, SBS7a and SBS7b both generated rwSNVs but affected distinct motif classes (**Figure S3a-b**). SBS7b preferentially disrupted proline-directed phosphorylation motifs targeted by CDK and MAPK kinases (X-[pS/pT]-P-X) (**Figure S2c**), whereas SBS7a was enriched in rwSNVs affecting protein glycosylation motifs and 14-3-3 binding sites involved in protein stability and scaffolding interactions ^32,33^. As expected, the rwSNVs from the two UV SBS signatures can be explained by the trinucleotide preferences of the signatures and corresponding substitutions in the genetic code (**Figure S3b**).

Together, these results demonstrate that distinct mutational signatures leave characteristic and mechanistically interpretable imprints on protein sequence motifs within cellular signaling networks.

### rwSNVs in cancer driver genes and pathways link mutational processes to functional impact

Cancer driver mutations confer oncogenic properties and are subject to positive selection in cancer genomes. To contextualise known cancer drivers and identify additional candidate genes, we asked whether driver mutations can be interpreted through the lens of mutational processes and rwSNVs. We therefore identified genes in which rwSNVs occurred more frequently than expected, given the background distribution of missense SNVs within individual cancer types and cohorts.

We identified 57 rwSNV-enriched genes across 15 cancer types (**Figure 3a, Table S4**; hypergeometric tests, FDR < 0.05), of which 20 are established cancer genes. These included major oncogenes and tumor suppressors such as *BRAF, CTNNB1, TP53, BCLAF1*, and *U2AF1*, some of which were independently enriched in multiple cancer types. Established cancer genes were significantly enriched among the rwSNV-enriched genes (hypergeometric test, P = 1.0 x 10^-5^), suggesting positive selection of rwSNVs in cancer.

**Figure 3.**
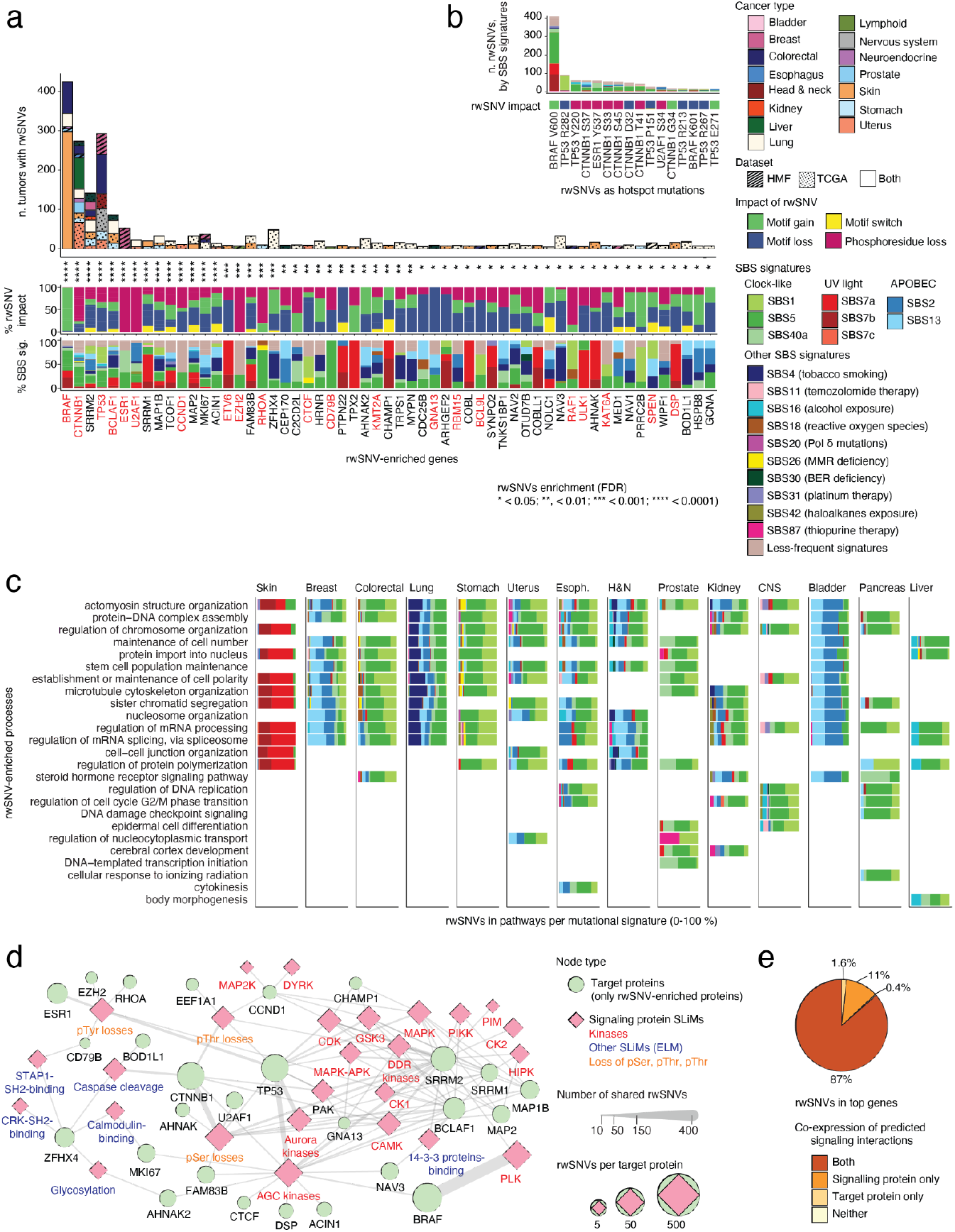
Cancer genes and pathways altered by rwSNVs and mutational processes. **(a)** Genes with significantly enriched rwSNVs (hypergeometric tests, FDR < 0.05). The bar plot (top) shows the number of cancer samples affected by rwSNVs in each gene. The color strips below show the functional impact of rwSNVs in the context of SLiMs (middle) and the mutational signatures associated with the rwSNVs (bottom). Established cancer genes are colored red. **(b)** rwSNVs representing the most frequent pan-cancer mutational hotspots (min. 10 cancer samples). Bar colors reflect SBS signatures associated with the rwSNVs while the color strip below shows the functional impact of rwSNVs on SLiMs. **(c)** Biological processes and pathways enriched in rwSNVs identified across multiple cancer types (FWER < 0.05 from ActivePathways). The relative contributions of SBS signatures to rwSNVs affecting genes in each pathway are shown as stacked bars, with colors indicating the associated mutational signatures. **(d)** Reconstruction of a putative signaling network involving interactions of rwSNV-enriched proteins and the signaling proteins binding rwSNV-altered SLiMs. Node labels are colored based on the type of signaling protein (kinases or other proteins) or rwSNV impact (phosphoresidue loss). **(e)** Co-expression analysis of rwSNV-enriched genes (*i*.*e*., target proteins) and the SLiM-associated signaling proteins based on matching transcriptomics data in TCGA. The pie chart shows the fractions of signaling interactions of signaling proteins and target proteins that are supported by gene expression in the cancer samples in which the relevant SLiMs and rwSNVs were identified. In panels (a-c), SBS signatures with at least two significant SLiM associations are listed and others are collapsed (shown in beige).

rwSNVs also encompassed well-characterized mutational hotspots, including BRAF V600E, TP53 R282W, and CTNNB1 S37F, and were recurrently observed in dozens to hundreds of cancer samples (**Figure 3b**). Integrating mutational signatures with SLiM annotations provided insight into the mutational processes driving these recurrent mutations, extending prior studies of phosphorylation-associated cancer mutations ^19,34^. For example, the rwSNVs TP53 Y220C and ESR1 Y537S caused loss of phosphotyrosine sites and were primarily attributed to clock-like mutational processes. rwSNVs in BCLAF1, a tumor suppressor and regulator of apoptosis ^35^, were associated with the tobacco-related signature SBS4 in lung cancer (**Figure S4a**), consistent with smoking-linked somatic alterations disrupting SLiMs in this cancer type. We also observed recurrent rwSNVs in CTNNB1 (β-catenin), where rwSNVs at position D32 in liver cancer were predicted to disrupt a caspase-cleavage motif from clock-like mutational signature SBS5, complementing the established role of D32 substitutions in β-catenin stabilisation ^21^ (**Figure S4b**).

To further assess the functional relevance of rwSNVs in these 57 prioritized genes, we analyzed variant impact scores using AlphaMissense ^36^ and CADD ^37^. rwSNVs in these genes showed significantly higher predicted impact than rwSNVs genome-wide, based on AlphaMissense scores (mean score 0.73 *vs* 0.43, P < 2.2 x 10^-16^; Wilcoxon test) with similar trends observed for CADD and mean of both scores (**Figure S5a-f**). The highest-impact rwSNVs frequently affected central oncogenes and tumor suppressors, including *TP53, ESR1*, and *CTNNB1*, consistent with their known roles in oncogenic pathways (**Figure S5g**).

To contextualize rwSNV functionally, we performed pathway enrichment analysis using ActivePathways ^38^, integrating rwSNV-enriched genes across cancer types. This analysis identified 158 significantly enriched biological processes and molecular pathways (Family-Wise Error Rate (FWER) < 0.05; **Figure 3c, Figure S6, Table S5**), including pathways central to cancer development such as cell-cycle regulation, chromatin remodeling, DNA damage response, and cell junction organization. rwSNVs within these pathways were driven by both ubiquitous and tissue-specific mutational processes, showing clock-like signatures across cancer types, APOBEC signatures in breast and bladder cancers, smoking-associated signatures in lung cancer, and UV-associated signatures in melanoma. These results indicate convergence of motif-rewiring mutations from distinct mutational processes onto shared signaling networks in cancer.

To further characterise the functional context of rwSNVs, we performed complementary network-based and transcriptomic analyses. We reconstructed a putative signaling network that comprised SLiM-specific protein-protein interactions involving the proteins encoded by the 57 rwSNV-enriched genes and their SLiM-specific signaling partners (**Figure 3d, Table S6a-b**). Network analysis highlighted central proteins with multiple motif-rewiring interactions, including CTNNB1, TP53, and BCLAF1, as well as splicing factors SRRM1 and SRRM2. rwSNVs in these frequently altered motifs bound by AGC kinases (PKA, PKC, AKT1) or proline-directed kinases (CDK, GSK3, MAPK), and often included phosphoserine losses. Analysis of matched TCGA transcriptomic data showed that 87% of predicted interacting protein pairs were co-expressed in corresponding cancer samples (**Figure 3e, Figure S1d-f**), supporting the plausibility of these interactions *in vivo*. We further examined this network using synthetic lethality data from the SynLethDB database ^39^. Among predicted interactions annotated in the database (15%), we identified eight synthetic lethal relationships, all involving TP53 and kinases interacting with rwSNV-altered SLiMs in TP53 (**Figure S7a-b**). Together, these analyses link rwSNVs in cancer genes to signaling network architecture and potential functional vulnerabilities.

### Mutational hotspots of motif rewiring in cancer genes

We examined the top rwSNV-enriched genes in detail. *BRAF* oncogene was the most significant rwSNV-enriched gene due to the recurrent, clinically actionable V600E hotspot mutation ^40,41^ observed in 425 samples, especially in melanoma (192 samples in TCGA; FDR = 4.3 x 10^-204^, one-tailed hypergeometric test; **Figure 4a-c**) as well as colorectal and lung cancers. BRAF V600E was predominantly associated with UV-related (SBS7b) or clock-like (SBS5) mutational signatures and was predicted to induce a phosphorylation motif bound by Polo-like kinase 1 (PLK1) (MIMP posterior p = 0.89). PLK1 regulates mitotic progression and DNA damage response and is a potential therapeutic target ^42^. While BRAF V600E constitutively activates MAPK/ERK signalling ^40^, our motif-rewiring analysis predicts an interaction with PLK1 as an additional signalling feature of this mutation. Consistent with this observation, recent studies have reported PLK1-dependent resistance to BRAF inhibition in melanoma and enhanced efficacy of BRAF-targeted therapy upon PLK1 co-inhibition in colorectal cancer models ^43,44^,

**Figure 4.**
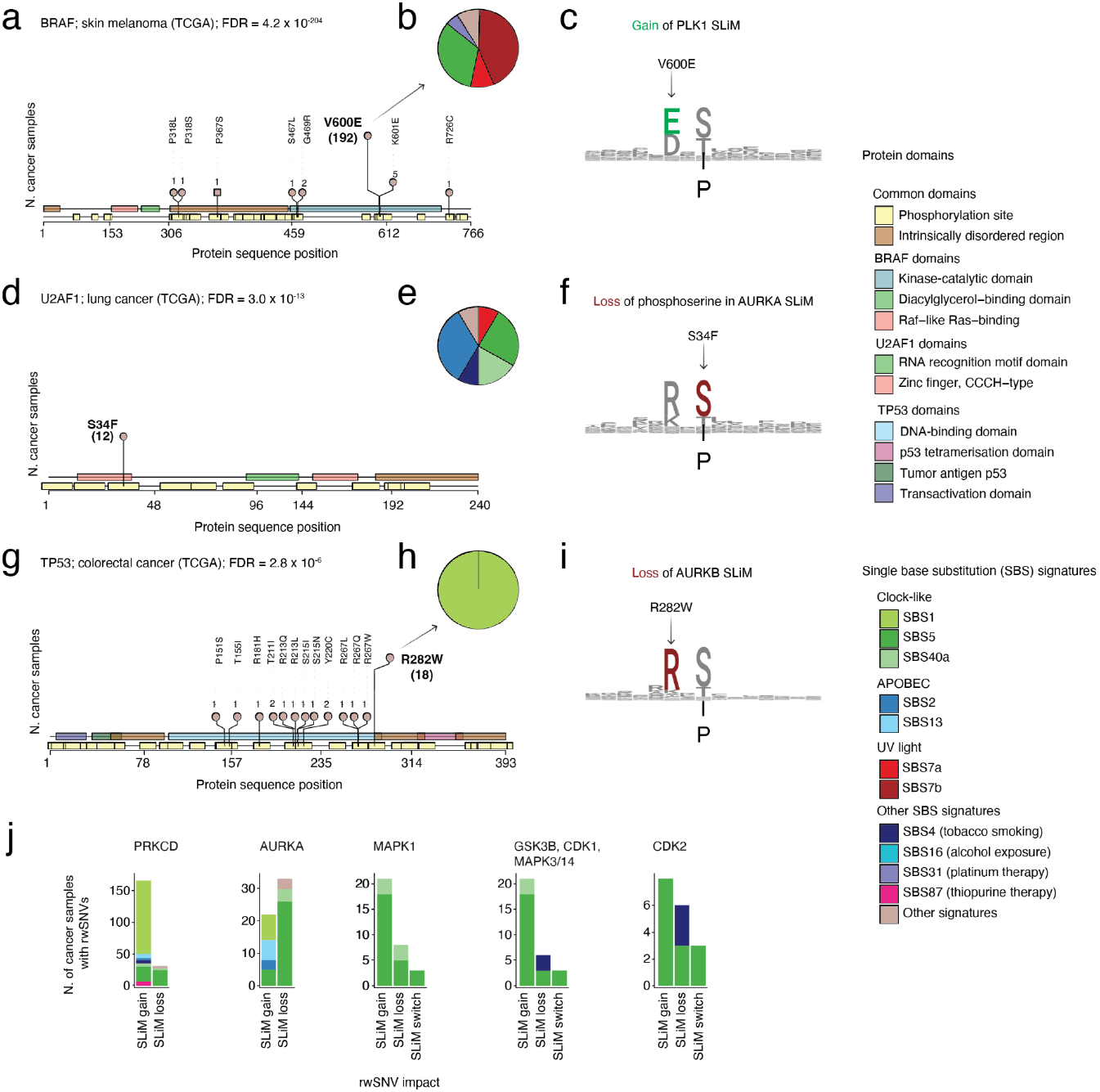
rwSNV hotspots in cancer driver genes are associated with specific mutational processes. **(a-c)** The rwSNV BRAF V600E in melanoma is often linked to UV-light signature SBS7b and induces a binding motif of Polo-like kinase 1. **(d-f)** The rwSNV U2AF1 S34F in lung cancer is linked to clock-like and APOBEC mutagenesis and removes a phosphoresidue in a binding site of Aurora kinase A. **(g-i)** The rwSNV TP53 R282W disrupts a SLiM of Aurora kinase B. Needle plots (left; panels a, d, g) show the locations of all rwSNVs in the protein sequences, with the highlighted rwSNVs shown in bold.. Protein domains and phosphorylation sites are shown as colored rectangles. Piecharts (middle; panels b, e, h) show the SBS mutational signatures contributing to the rwSNVs. SLiMs altered by the hotspot rwSNVs (right; panels c, f, i) are shown as sequence logos, with confirmed phosphoresidues (P) indicated. **(j)** Synthetic lethal interactions involving TP53 and the kinases binding the rwSNV-altered SLiMs. The barplot shows cancer samples having rwSNVs the SLiMs, grouped by functional impact of wrSNVs and mutational signatures (colors).

The splicing factor U2AF1 was significantly enriched in rwSNVs in lung cancer, driven by the recurrent S34F hotspot mutation identified in 12 lung cancer samples in TCGA (FDR = 2.8 x 10^-10^) (**Figure 4d-f**). This rwSNV substitutes a phosphorylated serine within the RNA recognition domain and was predicted by MIMP to disrupt an Aurora kinase-bound phosphosite, consistent with previous evidence linking Aurora signaling to splicing regulation ^45^. U2AF1 S34F substitutions were predominantly attributed to APOBEC-associated mutagenesis, a process frequently observed in aggressive lung cancers ^46^. Given the established role of U2AF1 in 3′ splice-site recognition ^47^, and the known impact of S34F on splice-site selection and RNA processing ^48^, these results link APOBEC-driven mutagenesis to motif-level disruption of post-transcriptional regulation in lung cancer.

The tumor suppressor TP53 was also enriched in rwSNVs across multiple cancer types, driven in part by the recurrent R282W rwSNV predominantly attributed to the clock-like SBS1 signature (*e*.*g*., primary colorectal cancer, FDR = 6.4 x 10^-13^) (**Figure 4g-i**). This rwSNV was predicted to disrupt SLiMs bound by Aurora kinase B (MIMP posterior p = 0.89) as well as ROCK, PKC and PKA kinases. Aurora kinases are involved in mitotic progression, chromosome segregation and cytokinesis ^49^.

Disruption of Aurora kinase B binding at R282W is supported by prior *in vitro* evidence ^20^. The R282W substitution occurs in the DNA-binding domain of TP53 and impairs transcriptional activation of target genes, thereby contributing to loss of tumor suppressor activity ^50^. To place these motif-rewiring effects in a broader functional context, we examined known genetic interaction data and identified a few reported synthetic lethal interactions between TP53 and kinases predicted to bind rwSNV-altered SLiMs (**Figure 4j, Figure S7a-b**). While these interactions were not systematically enriched across all rwSNVs, they illustrate how rwSNV-driven disruption of phosphorylation-dependent interactions may intersect with known genetic vulnerabilities. For example, the known synthetic lethal interaction between TP53 and PRKCD coincided with recurrent rwSNVs at position 282, which were predicted to disrupt a PRKCD binding site and were observed in 129 samples across 11 cancer types, predominantly associated with SBS1 (**Figure S7c**).

Together, these observations support the broader notion that linking mutational processes to motif-level network rewiring can provide mechanistic insights and highlight potential functional dependencies in cancer cells.

### Molecular drivers of APOBEC mutagenesis-induced rwSNVs

Lastly, we investigated molecular drivers of rwSNV formation, focusing on the endogenous APOBEC mutational process reflected in the signatures SBS2 and SBS13. APOBEC3 cytosine deaminases are involved in innate immunity ^51^ and are established as drivers of somatic mutagenesis ^52,53^. To associate rwSNVs with this mutational process, we correlated rwSNV burden attributed to SBS2 and SBS13 with median-dichotomized *APOBEC3A* expression in matched TCGA transcriptomes.

*APOBEC3A* expression was significantly associated with APOBEC-driven rwSNV burden in multiple cancer types (Kruskal-Wallis tests, FDR < 0.05; **Figure 5a, Table S7**). Samples with high *APOBEC3A* expression harbored significantly more rwSNVs from SBS2 and SBS13 than samples with low expression. The strongest associations were observed in lung adenocarcinoma (FDR = 1.6 x 10^-8^) and breast cancer (FDR = 3.8 x 10^-7^), with additional significant effects in uterine, head and neck, and lung squamous cell cancers. To refine these findings, we stratified rwSNVs by their functional impact on SLiMs and identified 74 significant associations with *APOBEC3A* expression (FDR < 0.05; **Figure 5b**). rwSNVs affecting SLiMs involved in caspase cleavage, calmodulin binding, SH2 and 14-3-3 domain interactions, and rwSNVs in kinase signaling motifs (PKA, AKT, CK2, PAK1) were associated with *APOBEC3A* expression across cancer types. Notably, rwSNV-driven losses of phosphoserine residues were highly correlated with *APOBEC3A* expression in multiple cancers, indicating widespread disruption of phosphorylation-dependent signaling by APOBEC mutagenic processes.

**Figure 5.**
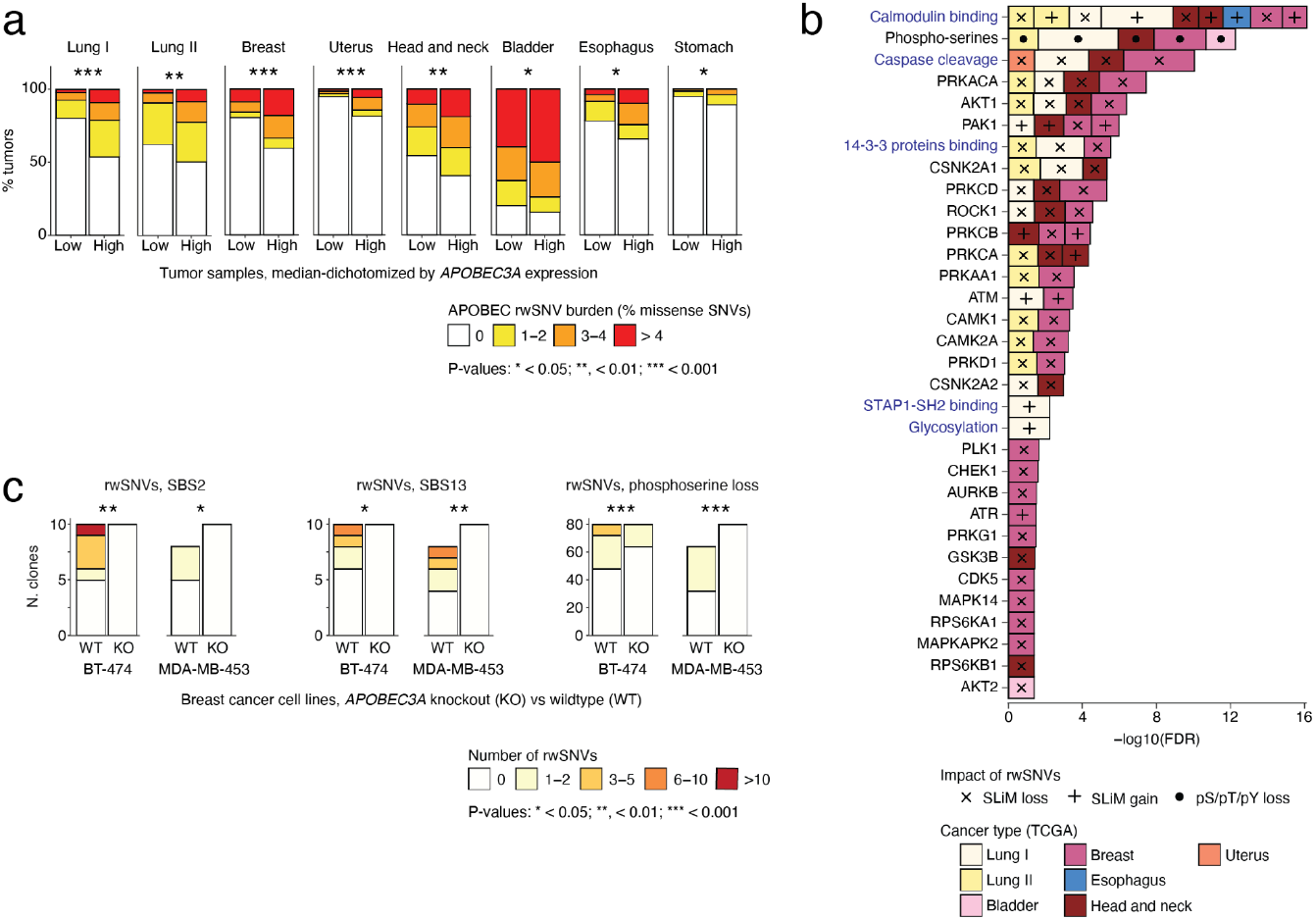
APOBEC-related rwSNVs in cancer genomes are associated with APOBEC3A activity. **(a)** Associations of rwSNV burden with *APOBEC3A* gene expression. Stacked barplots quantify rwSNV burden in cancer samples grouped by *APOBEC3A* gene expression, using matching cancer transcriptomes from TCGA. Cancer samples were median-dichotomized based on *APOBEC3A* expression (low vs. high). Colors show the fraction of rwSNVs in all missense SNVs, accounting for overall mutation burden in cancer samples. Each cancer type in TCGA was analyzed separately and significant results are reported (Kruskal-Wallis tests, FDR < 0.05). **(b)** Associations of *APOBEC3A* gene expression and rwSNV burden on individual SLiMs (Kruskal-Wallis tests, FDR < 0.05). Colors indicate cancer types, and shapes on bars reflect the functional impact of rwSNVs on SLiMs. SLiMs highlighted in blue are ELM SLiMs. Lung I: lung adenocarcinoma; Lung II: lung squamous cell carcinoma. **(c)** Comparison of rwSNV burden from APOBEC signatures SBS2 and SBS13 in breast cancer cell lines with *APOBEC3A* knockout and wildtype cell lines (derived from Petljak, *et al*., 2022 ^52^). P-values were computed using negative binomial regression (* < 0.05; **, < 0.01; *** < 0.001).

To obtain causal support for these associations, we analyzed whole-genome sequencing data from cancer cell lines with *APOBEC3A* knockout experiments ^52^. Breast cancer cell lines lacking *APOBEC3A* showed significantly fewer rwSNVs attributed to SBS2 and SBS13, as well as fewer rwSNVs causing phosphoserine losses, compared to wild-type controls (**Figure 5c, Table S8**). These results provide experimental support linking *APOBEC3A* activity to motif-rewiring mutations.

Together, these analyses demonstrate that endogenous APOBEC mutagenesis leaves measurable and functionally interpretable footprints in protein sequence motifs, linking molecular drivers of mutational processes to motif-rewiring alterations in cellular signaling networks.

## Discussion

Our study demonstrates that endogenous and environmental mutational processes leave distinct and mechanistically interpretable imprints on cancer signaling pathways. While mutational signatures are well established as descriptors of trinucleotide biases in cancer genomes ^3^, our analysis links these processes to specific amino acid substitutions that rewire SLiMs or post-translational modification sites within cellular signaling networks. SNVs altering these sequence motifs reflect mutational processes shaped by the genetic code, providing a direct connection between mutational processes and signaling-level functional consequences. Protein-coding substitutions arising from mutational processes leave distinct footprints on cellular signaling networks, such as UV light-induced disruptions of motifs bound by CDK or MAPK kinase motifs prominent in melanoma, or APOBEC-driven phosphoserine losses observed in multiple cancer types.

We propose that rwSNVs in SLiMs play dual roles in tumor evolution. A subset of rwSNVs undergo positive selection and contribute to oncogenesis through targeted disruption or creation of regulatory motifs, as evidenced by their enrichment in well-established cancer genes, and by the convergence of rwSNVs from distinct mutational processes onto shared signalling networks and hallmark cancer pathways. In contrast, many rwSNVs likely arise as by-products of genome-wide mutational processes and may alter other aspects of cellular signalling logic without directly driving oncogenesis, collectively contributing to inter- and intra-tumoral heterogeneity.

A key advantage of our framework is its ability to interpret missense SNVs located in IDRs, which may be less optimally captured by structure- or conservation-based variant effect predictors. By integrating mutational signatures with SLiM annotations, our approach provides a complementary strategy for interpreting the functional impact of missense mutations at signaling sites, linking the causes of somatic variation to their potential molecular consequences.

One illustrative example of the translational relevance of motif rewiring is the oncogenic BRAF V600E mutation. Beyond its canonical role in constitutive MAPK/ERK activation, our analysis suggests that BRAF V600E may introduce a *de novo* phosphorylation motif recognized by Polo-like kinase 1, thereby rewiring signaling interactions downstream of UV- or clock-like mutagenesis. This observation provides a potential mechanistic link between mutational etiology and therapeutic resistance. Clinically, resistance to targeted therapy remains a major challenge in BRAF V600E-mutant cancers, with activation of bypass pathways and adaptive signalling limiting long-term efficacy. Consistent with our motif-level predictions, recent preclinical studies show that targeting PLK1 can overcome resistance to BRAF inhibition ^43^ and that co-targeting PLK1 alongside MAPK pathway inhibitors produces synergistic anti-tumour effects ^44^, underscoring how perturbations in signalling networks highlighted by rwSNVs might identify actionable vulnerabilities. While experimental validation of our predictions is required, this example highlights how motif-level interpretation of driver mutations may reveal secondary signaling dependencies relevant to combination treatment strategies. More broadly, these findings showcase how environmental mutagens such as UV radiation can have context-dependent effects on human biology: while UV exposure is essential for vitamin D synthesis ^54^, it can also induce specific mutational processes that rewire signaling motifs and promote oncogenic pathways.

Our findings further demonstrate that mutational signatures of shared etiology can exert distinct functional effects on signalling networks in cancer. For example, the APOBEC-associated signatures SBS2 and SBS13, despite arising from related enzymatic activity, showed divergent impacts on SLiMs due to their different trinucleotide preferences. Similarly, UV-associated signatures SBS7a and SBS7b altered distinct classes of motifs. These observations underscore how subtle differences in mutational processes can translate into qualitatively different signaling perturbations, an effect that may be broadly relevant across cancer types.

Several limitations should be considered when interpreting our results. SLiMs represent only one layer of signaling regulation, and motif alteration alone is not sufficient to guarantee functional interaction changes, which also depend on protein expression, localisation, and post-translational context. In addition, SLiMs are short and only depend on a few core residues, leading to unavoidable false-positive predictions. Our analysis is further limited to well-characterized mutational signatures and is better powered for common signatures and motifs. Nevertheless, the major associations we identify are reproducible across independent cohorts and supported by experimental phosphosite annotations and APOBEC knockout data, lending confidence to the robustness of our conclusions.

In summary, our study links environmental and endogenous mutational processes to motif-level rewiring of cellular signaling networks. By connecting mutational etiology to proteomic and signaling consequences, this framework provides a complementary method for interpreting missense variation in cancer and may inform future efforts to identify biomarkers, signaling vulnerabilities, and context-specific therapeutic strategies.

## Methods

### Cancer genomics datasets.

We analyzed somatic single nucleotide variants (SNVs) from two cancer genomics cohorts: whole-exome sequencing (WES) data of primary cancers from the Cancer Genome Atlas (TCGA) Pan-Cancer Atlas project ^26^, and whole-genome sequencing (WGS) data of metastatic cancers from the Hartwig Medical Foundation (HMF) project ^27^. As part of previous research efforts, DNA sequencing of tumor tissues was performed by members of the TCGA and HMF consortia, under approved Institutional Review Board protocols and along with written informed consent of patients. This study is covered by the University of Toronto Research Ethics Board protocol 37521. For TCGA, we used the publicly available Multi-Center Mutation Calling in Multiple Cancers (MC3) dataset ^55^ and retained the SNVs that passed the MC3 quality filter, and excluded hypermutated samples (>1800 variants for WES data). For HMF, we also used previously preprocessed data, excluded hypermutated samples (>90,000 variants for WGS data), and excluded SNVs that did not pass the HMF variant selection filters. We also removed 288 duplicate samples from tumors of the same patients in HMF by selecting samples with the highest tumor purity. We also excluded samples with low signature reconstruction accuracy (see below). For further stringency, we excluded variants with high frequency in the human population (allelic frequency > 0.05) based on the Genome Aggregation Database (gnomAD) ^56^ (v.4.1). To jointly analyze TCGA and HMF data, we considered 18 consolidated cancer types. Protein-coding impact of SNVs was derived using the ANNOVAR software ^57^ (v.2022-08-02) based on human genome version GRCh37 and canonical protein isoforms as listed in ActiveDriverDB ^17^. In total, we extracted and analyzed 1,240,612 missense SNVs.

### Mutational signatures.

We used the SigProfilerAssignmentR ^28^ method (v. 0.0.17) to assign COSMIC v3.4 ^58^ mutational signatures to SNVs, with TCGA data analyzed in WES mode and HMF data in WGS mode using sample-level VCF files as input and default parameters for signature decomposition. Exposure profiles for each sample and reconstruction similarities of observed and predicted mutational signatures were derived for each cancer genome, followed by quality assessment via cosine similarity of reconstructed and observed mutational profiles. Samples with reconstruction accuracy ≥70% were retained.

### Predicting SNV impact on SLiMs in phosphosites.

We used the MIMP machine learning method ^20^ to predict the effect of missense SNVs on short linear motifs (SLiMs) based on position weight matrices (PWMs). 124 kinase-bound SLiMs of length 15 (*i*.*e*., phosphoresidue with ±7 residues) developed previously in the MIMP study ^20^ were mapped to experimentally detected phosphosites from the ActiveDriverDB database (v2021) ^17^, which aggregated data from multiple databases (PhosphoSitePlus ^59^, UniProt ^60^, Phospho.ELM ^61^, HPRD ^62^). To define rwSNVs, missense SNVs associated with SLiM gains or losses were selected based on a posterior probability from MIMP (p > 0.8). If the same rwSNV led to loss of one SLiM and gain of another SLiM, it was annotated as a motif switch. In addition, we selected missense SNVs substituting phosphoresidues (serines, threonines, tyrosines).

### Predicting SNV impact on SLiMs from ELM.

We analyzed SNVs using additional SLiMs from the Eukaryotic Linear Motif (ELM) database ^63^ (v.2023-01). Positive sequence instances of SLiMs were extracted from the ELM database, excluding the ones with missing residues and proximal to protein sequence termini. We selected human or mouse SLiMs, excluded SLiMs with few positive sequences (<20), and further excluded SLiMs that were redundant with kinase-bound SLiMs from the MIMP dataset. ELM SLiMs were mapped to intrinsically disordered regions (IDR) of proteins using ActiveDriverDB. We updated the MIMP method to permit analysis of ELM SLiMs. SLiMs were extended to length 15 for consistency with kinase-bound SLiMs at phosphosites (described above). Then we aligned sequence instances using the R package muscle ^*64*^, ensuring no alignment gaps (gapopen = 1e-20) and otherwise default settings. From each alignment, a 15-residue window centered on the core motif was extracted to compile the positive sequences. For negative sequences, we randomly sampled 10,000 sequences of length 15 from IDRs.

Gaussian mixture models in MIMP were adapted to ELM motifs. For each SLiM, we modeled sequence specificity using PWMs constructed from positive sequences following the method described in the MIMP study ^20^, with four modifications: (i) at least 20 positive sequences were required for PWM construction; (ii) no prior probability was used for the positive distribution (*i*.*e*., prior p = 1), (iii), central sequence positions were included in the motif matching scores, and (iv) an additional step of iterative refinement of positive sequences was performed to exclude the lowest-scoring positive sequences based on Tukey’s outlier method ^65^. In total, 20 ELM SLiMs were included in the analysis (CLV_C14_Caspase3-7, DOC_CYCLIN_RxL_1, DOC_MAPK_JIP1_4, LIG_14-3-3_CanoR_1, LIG_AP2alpha_2, LIG_CaM_IQ_9, LIG_CtBP_PxDLS_1, LIG_EH_1, LIG_LIR_Gen_1, LIG_NRBOX, LIG_SH2_CRK, LIG_SH2_GRB2like, LIG_SH2_STAP1, LIG_SH3_CIN85_PxpxPR_1, LIG_TRAF6_MATH_1, MOD_CMANNOS, MOD_GSK3_1, MOD_N-GLC_1, MOD_SUMO_for_1, TRG_ER_FFAT_1).

### Associating rwSNV burden with mutational signatures.

To associate rwSNVs with mutational signatures, we analyzed the number of rwSNVs associated with each signature relative to the signature’s contribution to all missense SNVs using one-tailed hypergeometric tests. We excluded artefact SBS signatures according to COSMIC ^31^ (SBS27, SBS43, SBS45, SBS46, SBS47, SBS48, SBS49, SBS50, SBS51, SBS52, SBS53, SBS54, SBS55, SBS56 SBS57, SBS58, SBS59, SBS60, SBS95). Associations were computed separately for each of the cancer types and cohorts (TCGA, HMF) and rwSNV impact (loss of phosphoserine -threonine or-tyrosine, alterations of phosphosite SLiMs, and alterations of SLiMs from ELM. P-values were adjusted for multiple testing using the Benjamini-Hochberg FDR method ^66^ and significant results were selected (FDR < 0.05). We extended this analysis to each type of SLiM using one-tailed hypergeometric tests, tested each cancer type and cohort separately, adjusted P-values for multiple testing, and selected significant results (FDR < 0.05). This covered 124 kinase-bound SLiMs, 20 additional SLiMs from ELM, and known phosphoresidue substitutions (experimentally confirmed phosphoserines, -threonines, or -tyrosines). Next, rwSNVs and mutational signatures were interpreted using network analysis via the Cytoscape software ^67^. We clustered SLiMs into subnetworks based on rwSNVs, assuming that the same rwSNVs often affected similar SLiMs. Nodes in this SLiM enrichment network were visualized as pie charts that reflected the proportions of rwSNVs associated with mutational signatures and node sizes reflected log-scale rwSNV counts.

### Gene expression analysis of signaling interactions involving rwSNVs.

We used matching transcriptomics data from TCGA to investigate functionality of rwSNVs. We extracted mRNA expression levels quantified as fragments per kilobase of transcript per million mapped reads upper quartile (FPKM-UQ) for (i) genes harboring rwSNVs (encoding target proteins having rwSNV-altered SLiMs) and (ii) genes encoding upstream signaling proteins binding the SLiMs. This analysis only included the SLiMs impacted by rwSNVs: 76 kinases from the MIMP dataset and additional interactions involving signaling proteins corresponding to 8 ELM SLiMs (**Table S9**). For each rwSNV and its associated interaction of signaling protein and target protein, the mRNA expression status was evaluated per cancer sample having a specific rwSNV and considering the mRNA level of target protein and also the mRNA level of the rwSNV- and SLiM-associated signaling protein. If multiple signaling proteins were associated with the same SLiM and rwSNV, the protein with the highest mRNA-based expression was selected. rwSNVs were then categorized according to whether (1) both the rwSNV-altered gene (*i*.*e*., the target protein) and the corresponding signaling protein were expressed, (2) only the mutated gene (*i*.*e*., the target protein) was expressed, (3) only the corresponding signaling protein was expressed, or (4) neither protein was expressed at the mRNA level. This analysis integrated matching transcriptomics data for nearly all rwSNVs found in genomics profiles in TCGA; however, for a few rwSNVs with multiple transcriptomics profiles available per cancer sample, mean gene expression values across the samples were used.

### Enrichment of rwSNVs in individual genes.

To identify positive selection in individual genes in cancer genomes, we analyzed rwSNVs in each protein-coding gene using one-tailed hypergeometric tests, considering the distribution of missense SNVs in the gene of interest relative to all protein-coding genes combined. Each cancer type and cohort was analyzed separately, genes with fewer than five rwSNVs in cancer samples were excluded from the analyses, the resulting P-values were adjusted for multiple testing using FDR, and significant genes were selected (FDR < 0.05). Known cancer genes from the COSMIC Cancer Gene Census ^58^ and IntOGen ^68^ databases were evaluated relative to all protein-coding genes using a hypergeometric test. Missense SNVs were visualized using the lolliplot method of the trackViewer R package ^69^. Protein domain information was retrieved from the Pfam database^70^. Functional impact of rwSNVs was evaluated using CADD ^37^ and AlphaMissense ^36^ scores derived from ANNOVAR. Lastly, we visualized rwSNV-enriched genes as a network of interactions comprising target proteins and signaling proteins, by selecting the interactions involving signaling proteins of rwSNV-altered SLiMs. rwSNVs involving phosphoresidue substitutions were also added to the network as meta-nodes. The network was visualized using the Cytoscape software ^67^.

### Pathway analysis of rwSNVs

Pathway enrichment analysis was performed using the ActivePathways method ^38^ by integrating gene-based rwSNV enrichment signals across multiple cancer types. As input, we used a matrix of gene P-values derived above with columns representing cancer types. To consolidate gene-based enrichments from TCGA and HMF, the most significant P-value for each gene was selected from the two cohorts. Gene sets representing biological processes of Gene Ontology and molecular pathways of the Reactome database ^71^ were derived from the g:Profiler web server ^72^ (downloaded 23 June 2024). Gene sets with 100 to 500 genes were analyzed and significant enrichments were selected (ActivePathways, with Holm family-wise error rate (FWER) correction; FWER < 0.05). Results were visualized as an enrichment map ^73^ using Cytoscape ^67^, followed by manual curation of subnetworks to display common functional themes for each sub-network.

### Analysis of synthetic lethality interactions involving rwSNVs.

To obtain functional evidence of rwSNVs, we examined the reconstructed signaling network of rwSNV-enriched genes and SLiM-mediated interactions for evidence of synthetic lethality. We examined the SynLeth Database ^74^ for five types of interactions: synthetic lethal (SL), non-synthetic lethal (NON-SL), synthetic dosage lethal (SDL), synthetic rescue (SR), and synthetic dosage rescue (SDR). We assigned each interaction from our dataset into a single category based on a weighted quantitative score, following the approach described previously ^39,74^, both including and excluding computationally predicted pairs. We selected the rwSNV-mediated gene pairs detected in the database (all involving *TP53*) as synthetic lethal and further analyzed these to identify the SLiMs, rwSNVs, and mutational processes.

### Clinical and molecular associations of rwSNVs.

To detect causal associations between mutational processes and rwSNV burden, we focused on APOBEC mutagenesis and analyzed matching transcriptomes from TCGA separately for each cancer type. *APOBEC3A* expression levels (transcripts per million) were median-dichotomized (high *vs*. low expression). Associations with *APOBEC3A* expression were analyzed in the context of rwSNV burden in two ways: first, for all rwSNVs combined, and second, separately for each type of SLiM. rwSNV burden was defined as the fraction of rwSNV count over total missense SNV count in the given cancer cohort to account for variations in overall mutation burden. rwSNV burdens between *APOBEC3A*-high -and *APOBEC3A*-low cancer samples were compared using Kruskal-Wallis tests, P-values were adjusted for multiple testing and significant results were selected (FDR < 0.05).

### Quantifying rwSNVs in *APOBEC3* gene knockout experiments.

To associate APOBEC mutational processes with rwSNV burden, we analyzed WGS data from previously generated isogenic cancer cell lines with *APOBEC3A* knockout (KO) ^52^. SNVs from *APOBEC3A* wild-type (WT) and *APOBEC3A* KO treatment groups for only the breast cancer cell lines were analyzed (breast cell lines: BT-474, MDA-MB-453). Clones flagged as non-unique in the original study were excluded, and only the daughter clones were analyzed. Functional impact of SNVs on protein-coding genes was annotated using ANNOVAR and mutational signatures were assigned using SigProfilerAssignment with COSMIC v3.4 SBS signatures, as described above. APOBEC signatures assigned to missense SNVs were selected and rwSNVs were determined using MIMP as described above. For each clone within each cell line, we counted total rwSNVs, rwSNVs associated with APOBEC mutational signatures (SBS2 or SBS13), and rwSNVs causing phosphoserine losses. To compare rwSNV burden in the context of APOBEC3 gene knockouts, we fitted negative binomial regression models to clone-level counts, by testing treatment group (KO vs WT) as a covariate in the alternative model, and compared them to intercept-only models using likelihood-ratio chi-square tests. Each cell line and each type of rwSNV (SBS2, SBS13, phosphoserine) was analyzed separately. Effect sizes were summarized as log2 fold-changes in rwSNV burden between KO and WT clones.

## Supplementary tables

**Table S1. Motif-rewiring msSNVs (rwSNVs)**. rwSNVs found in the TCGA dataset are shown. For each rwSNV, we report the associated SBS mutational signature and the predicted functional impact of SLiMs, based on the MIMP method (posterior p > 0.8). Since rwSNVs can impact multiple SLiMs, rows represent pairs of rwSNVs and SLiM instances rather than unique rwSNVs. SLiM type is reported as NA for rwSNVs corresponding to losses of phosphoresidues (serines, threonines, tyrosines). rwSNV results from the Hartwig Medical Foundation (HMF) dataset are not shown due to data access restrictions.

**Table S2. Mutational signatures enriched in rwSNVs**. SBS mutational signatures that were significantly enriched in rwSNVs in each cancer type and cohort (one-tailed hypergeometric tests). P-values were adjusted for multiple testing across the signatures using the Benjamini-Hochberg False Discovery Rate (FDR) method and filtered for significance (FDR < 0.05).

**Table S3. Mutational signatures enriched in rwSNVs in different classes of SLiMs**. SBS mutational signatures that were significantly enriched in rwSNVs in different classes of SLiMs in each cancer type and cohort (one-tailed hypergeometric tests). P-values were adjusted for multiple testing within each cohort using FDR and filtered for significance (FDR < 0.05).

**Table S4. rwSNV-enriched genes**. Genes having significant enrichments of rwSNVs relative to other protein-coding genes and all missense SNVs are shown (one-tailed hypergeometric tests). Each cancer type and cohort was analysed separately. P-values were adjusted for multiple testing within each cohort using FDR and filtered for significance (FDR < 0.05).

**Table S5. rwSNV-enriched biological processes and molecular pathways**. Molecular pathways and biological pathways enriched in rwSNVs in multiple cancer types, as identified by the ActivePathways method (FWER < 0.05). P-values from pathway analysis were adjusted for multiple testing using the Holm family-wise error rate (FWER) method as per default settings of ActivePathways. For each significant pathway, the overlapping genes having rwSNV enrichments are provided as Ensembl IDs.

**Table S6a. Reconstructed network of SLiM-specific interactions rwSNV-enriched target proteins and signaling proteins**. Each row includes an interaction between one of rwSNV-enriched genes and a signaling protein predicted to bind an rwSNV-altered SLiM in the target protein (as encoded by the rwSNV-enriched gene). Signaling proteins were annotated by using a kinase family-based grouping in the network visualization (Table S6b), while additional signaling proteins corresponding to ELM SLiMs or rwSNVs causing phosphoresidue losses are shown separately.

**Table S6b. Mapping of kinase families and kinases for the reconstructed signalling network**. Individual kinases mapped kinase families as used in the reconstructed network involving rwSNV-enriched genes and SLiM-specific interactions with signaling proteins (Table S6a).

**Table S7. Associating APOBEC-driven rwSNVs with *APOBEC3A* expression in cancer samples**. Primary cancer samples from TCGA were analyzed for APOBEC-associated rwSNVs (SBS2/SBS13) and median-dichotomized *APOBEC3A* expression from matching tumor transcriptomes. rwSNV burden was normalized relative to all missense SNVs and this relative rwSNV burden was compared in *APOBEC3A*-high vs *APOBEC3A*-low cancer samples using Kruskal-Wallis tests separately for each cancer type in two sets of comparisons, firstly across all functional types of SLiMs (denoted as Total Effect), and secondly, separately for each class of SLiM. P-values were adjusted for multiple testing for the two sets of comparisons using FDR and filtered for significance (FDR < 0.05). Lung I: lung adenocarcinoma; Lung II: lung squamous cell carcinoma.

**Table S8. Analysis of APOBEC-driven rwSNVs with *APOBEC3A* knockout in cancer cell lines**. rwSNVs were mapped from whole-genome sequencing data from *APOBEC3A* knockout (KO) and wildtype (WT) cancer cell lines from Petljak *et al*. (2022). rwSNV counts from APOBEC signatures (SBS2, SBS13) in clones were analyzed using negative binomial regression and P-values from chi-square tests are reported along with rwSNV counts.

**Table S9. Kinases and other signalling proteins used in the gene expression analysis**. Gene symbols of kinases and other signalling proteins interacting with SLiMs in rwSNV-enriched genes in the reconstructed signalling network, used to quantify gene expression in cancer samples in TCGA in which the rwSNVs were identified. Only SLiMs altered by at least one rwSNV in our analysis are included.

## Data availability.

Processed data are available as supplementary datasets. Input datasets from Hartwig Medical Foundation (HMF) representing WGS, transcriptomics, and metadata annotations of cancer samples of multiple cancer types are controlled-access and can be provided by HMF pending scientific review and a completed material transfer agreement. Requests for these datasets should be submitted to the Hartwig Medical Foundation. Input datasets from the Cancer Genome Atlas (TCGA) study representing WES, transcriptomics, and metadata annotations of cancer samples of multiple cancer types are available from the Genomic Data Commons website. Intermediate data files cannot be shared due to the use of controlled-access datasets from HMF.

## Acknowledgments

We would like to thank Drs. Sagi Abelson, Norman Davey, Ylva Ivarsson, and Hannes Röst for valuable discussions. This work was partially supported by the Canadian Institutes of Health Research (CIHR) Project Grants (PJT-162410, PJT-197925) and Catalyst Grant (DV1-197665) to J.R., the Investigator Award to J.R. from the Ontario Institute for Cancer Research (OICR), Discovery Grant of the Natural Sciences and Engineering Research Council (NSERC) (RGPIN-2023-04646) to J.R., and the New Investigator Award of the Terry Fox Research Institute (TFRI-PROJECT-1095) to J.R. A.B. was supported by the Ontario Graduate Scholarship (OGS). M.B. and K.C. were partially supported by fellowships from the Medical Biophysics Department at University of Toronto. M.B. was supported by a Caven Fellowship. M.F. was supported by the Spanish State Research Agency (Agencia Estatal de Investigación, AEI) grant FPI PRE2021-101048. Funding to OICR is provided by the Government of Ontario. The results shown here are in whole or part based upon data generated by the TCGA Research Network: https://www.cancer.gov/tcga. This publication and the underlying study have been made possible partly based on the data that Hartwig Medical Foundation has made available to the study.

## Author Contributions.

JM. and M.B. led data analyses. J.M. led data visualisation and interpretation. J.M. and J.R. wrote the original draft of the manuscript. J.M., M.B., N.A., K.C., A.B. Z.K., and J.R. reviewed and edited the manuscript. J.R. conceptualized the idea, supervised the project, and acquired funding. J.M., M.B., M.F., N.A., K.C., Z.K., and J.R. contributed to data analyses, interpretation and visualization. J.M., M.B., M.F., N.A., K.C., A.B., O.W., and J.R. contributed to methodology. All coauthors reviewed and approved the final manuscript.

