## Supplementary Figures for "Mutational signatures in cancer genomes alter protein sequence motifs in cellular signaling networks"

### Supplementary materials

J. Mishra, M. Bayati *et al*. (2026)

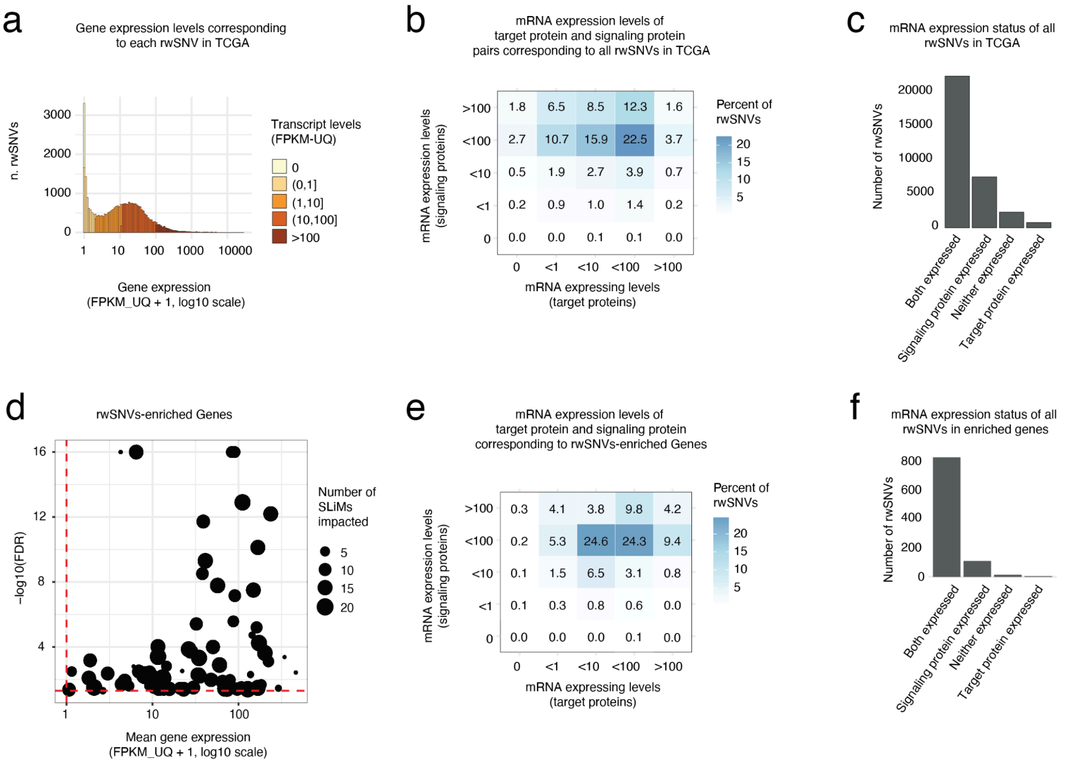


**Figure S1. Co-expression of rwSNV-affected target proteins and their SLiM-associated signalling partners in TCGA.** Using matched TCGA RNA-seq data, we quantified gene expression for genes involved in rwSNVs and SLiM-mediated protein-protein interactions with signalling proteins. **(a)** Expression levels of genes harboring rwSNVs across cancer cohorts. **(b)** All rwSNVs were binned by discretized mRNA expression of the target gene (i.e., the rwSNV-bearing gene) and its corresponding signaling protein binding the related SLiM. The proportion of rwSNVs falling into each expression-based bin is shown (color scale). **(c)** Corresponding rwSNV counts are shown for four expression states: target gene expressed only, signaling partner expressed only, both genes expressed, or neither gene expressed. **(d)** For genes significantly enriched in rwSNVs (FDR < 0.05, with at least 5 patients per rwSNV across each cancer cohort), the scatter plot shows gene-level rwSNV enrichment versus mean mRNA expression in the corresponding TCGA cohort. Point size denotes the number of distinct SLiMs rewired by rwSNVs in each target protein-coding gene. The horizontal dashed red line indicates the rwSNV enrichment significance threshold (FDR = 0.05), while the vertical dashed red line marks the mRNA expression detection threshold (FPKM-UQ = 1). **(e)** Same as panel (b), limited to the subset of rwSNVs occurring in the significantly rwSNV-enriched genes prioritized from multiple cancer types. **(f)** Same as panel (c), showing absolute rwSNV counts across the four expression states for the rwSNV-enriched gene set only.

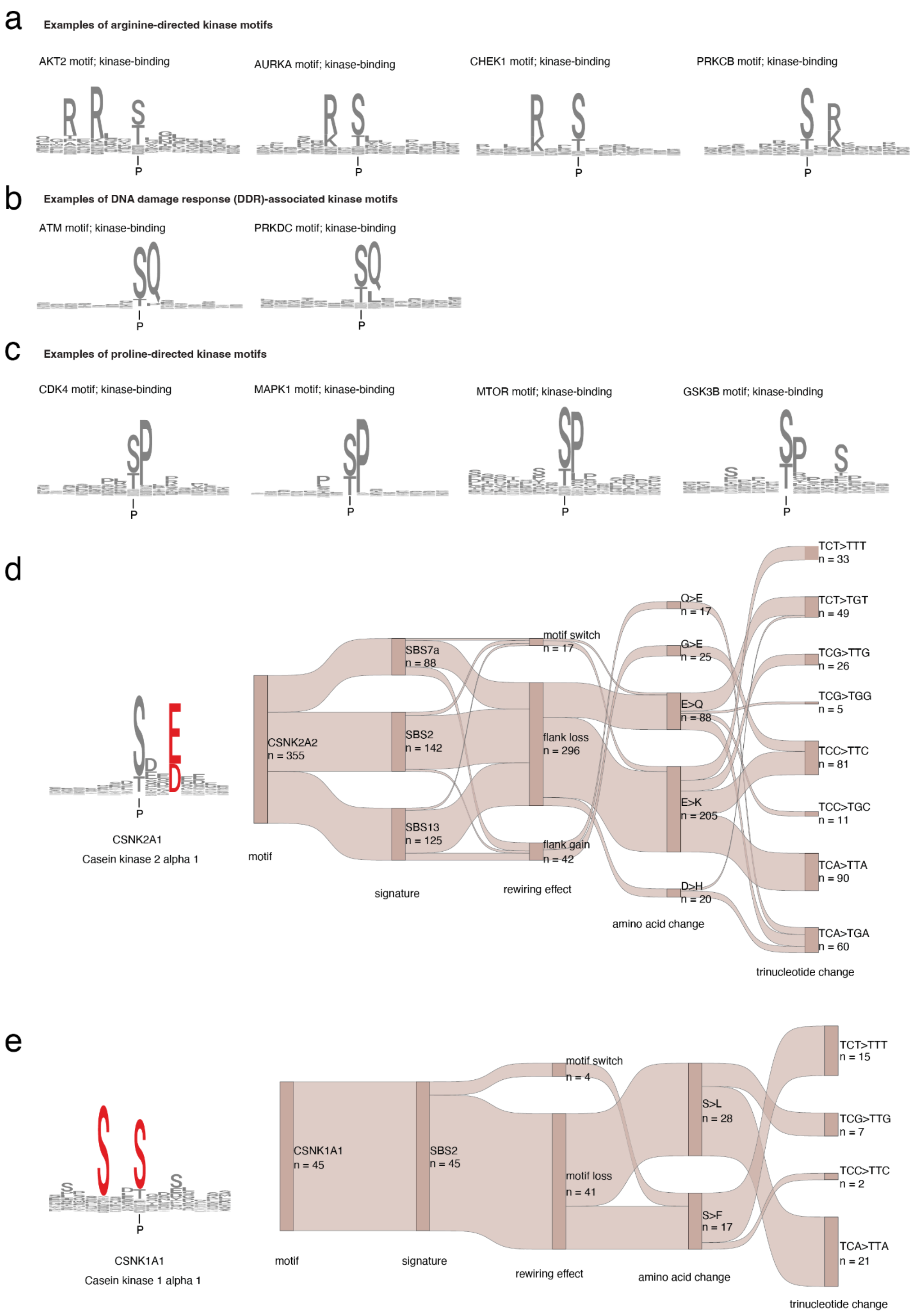


**Figure S2. Different classes of kinase-bound SLiMs and mutational signatures of rwSNVs altering the SLiMs of CK1 and CK2 kinases. (a-c)** Motif logos of representative phosphorylation motifs: **(a)** arginine-directed kinase motifs, **(b)** proline-directed kinase motifs, and **(c)** DNA damage response-associated kinase motifs. **(d)** rwSNVs affecting the SLiMs bound by CSNK2A1 (CK2) are shown together with the contributing trinucleotide contexts of SBS signatures and the related amino-acid substitutions based on the genetic code. The CSNK2A1 motif logo is shown on the left. **(e)** Same as panel (d), shown for CSNK1A1 (CK1)-associated SLiMs. Phosphorylated residues in the logos are indicated with the letter P.

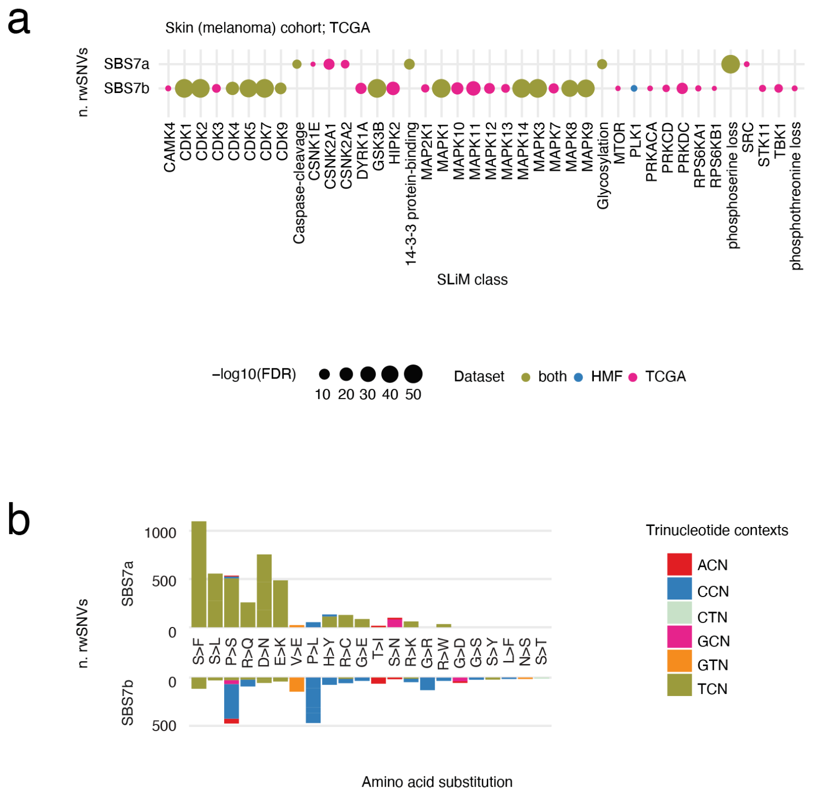


**Figure S3. Trinucleotide contexts and amino-acid substitutions of UV-light signatures SBS7a and SBS7b in melanoma.** Distinct motif-rewiring associations of SBS7a and SBS7b are explained by their trinucleotide preferences and amino-acid substitution biases. SBS7a is enriched in T[C>T]N nucleotide substitutions often resulting in S>F and S>L amino acid substitutions, while SBS7b is enriched in T[C>T]N nucleotide substitutions often resulting in P>L and P>S amino acid substitutions. This potentially reflects different UV-induced DNA lesions, cyclobutane pyrimidine dimers versus 6–4 photoproducts [1]. (a) Associations of mutational signatures and rwSNVs affecting SLiMs in melanoma, shown for TCGA, HMF, and both cohorts combined. (b) Comparison of frequency distributions of rwSNV-induced amino-acid substitutions attributed to SBS7a and SBS7b, annotated by underlying trinucleotide changes, highlighting how context-dependent biases may underlie the distinct motif-rewiring effects observed for SBS7a- versus SBS7b-driven rwSNVs.

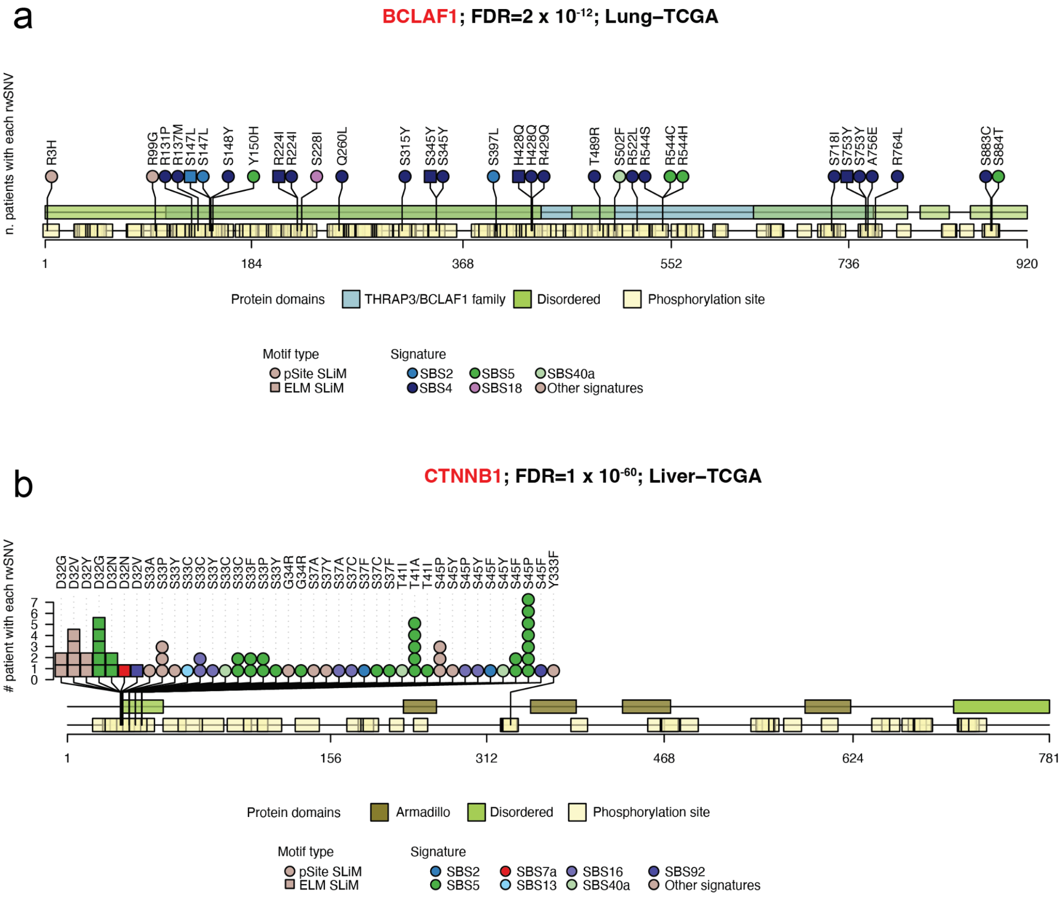


**Figure S4. enriched rwSNVs in BCLAF1 and CTNNB1.** Lollipop plots show the positions of rwSNVs along the protein sequence encoded by genes **(a)** *BCLAF1* and **(b)** *CTNNB1*. Protein domains and phosphorylation sites are shown as coloured rectangles. Each shape (circle or square) represents an rwSNV; color denotes the associated mutational signature, and shape indicates the type of SLiM. **(a)** rwSNVs in tumor suppressor BCLAF1 (BCL2 associated transcription factor 1) were mostly attributed to the tobacco-associated signature SBS4 in the lung cancer cohort of TCGA, supporting a smoking-linked disruption of regulatory motifs. **(b)** Recurrent rwSNVs in CTNNB1 (beta-catenin) were mostly attributed to the clock-like signature SBS5, as well as other signatures in the liver cancer cohort in TCGA. rwSNVs as position D32 were predicted to disrupt a caspase-cleavage motif. While D32 substitutions have previously been linked to β-catenin stabilisation [2], loss of a caspase cleavage site suggests a complementary mechanism of β-catenin accumulation and Wnt pathway activation. In addition, multiple rwSNVs in CTNNB1, arising from the clock-like signature SBS5 and the alcohol-linked signature SBS16, led to phosphoserine losses.

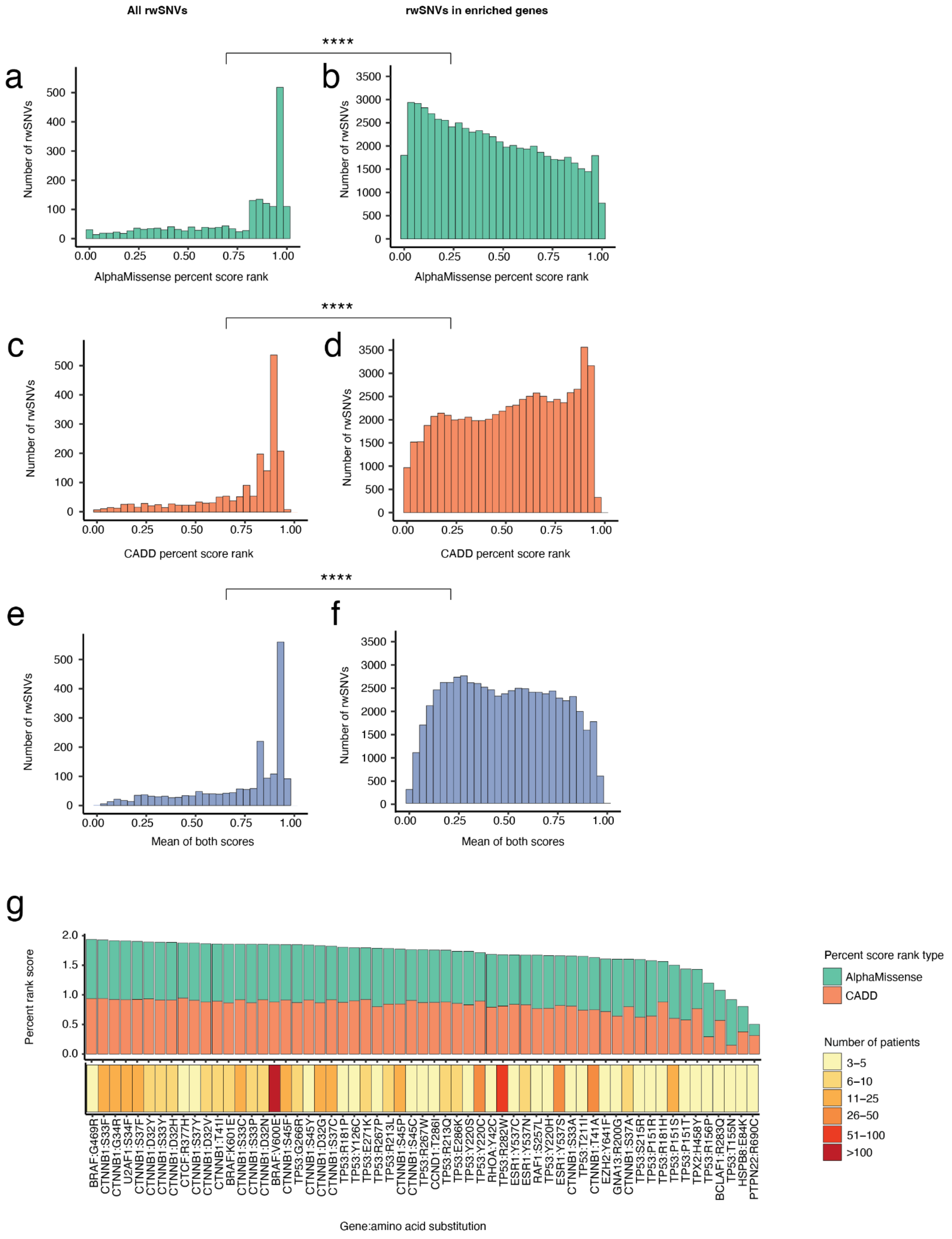


**Figure S5. Variant impact scores of rwSNVs.** Panels **(a,d)** show percentile ranks of variant impact scores from AlphaMissense, **(b,e)** show percentile ranks of impact scores from CADD, and **(c,f)** show the mean ranks of the two types of scores. Scores for all rwSNVs **(a, c, e)** and for rwSNVs in rwSNV-enriched genes **(b, d, f)** are compared. P-values were computed using one-tailed Wilcoxon rank-sum tests separately for each score type. Asterisks indicate statistical significance (****, $P < 2.2 \times 10^{-16}$). **(g)** Variant impact scores for the most recurrent rwSNVs observed in the cancer samples across the two cohorts. Barplots show variant impact scores of the rwSNVs (top) and the color strip indicates the number of cancer samples with the rwSNVs (bottom). rwSNVs are ordered by patient recurrence and the combined score. rwSNVs found in two or more cancer samples are shown.

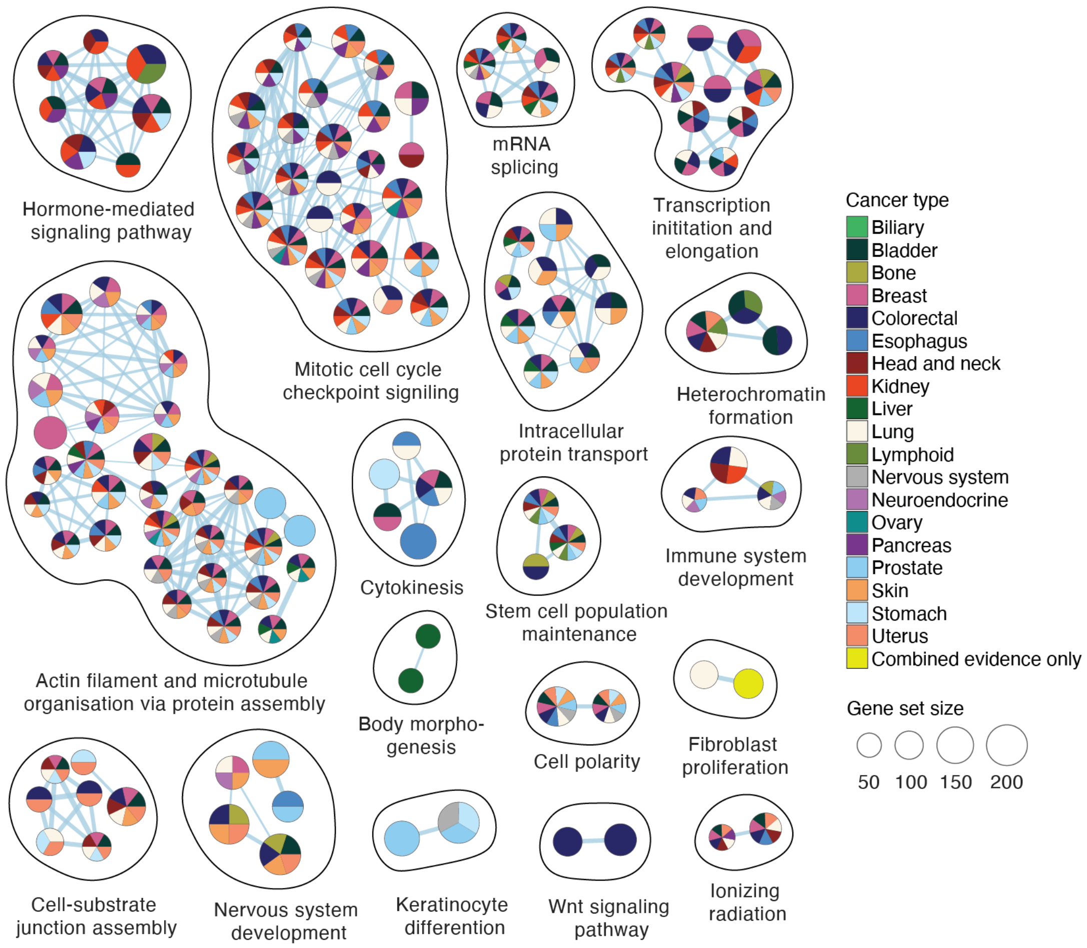


**Figure S6. Pathway enrichment analysis of rwSNV-enriched genes.** rwSNV-enriched pathways were identified by integrating genes with rwSNV enrichments across different cancer types using the ActivePathways method. For each gene, the most significant P-value from the TCGA or HMF dataset was selected for integration. Enriched pathways were selected based on significance (FWER < 0.05) and visualized as an enrichment map, a network of associations that displays enriched pathways as nodes, with edges connecting pathways sharing multiple genes. Node colors indicate the cancer type in which the corresponding rwSNV-based enrichment was detected. Yellow nodes indicate pathways that were found only when rwSNVs from multiple cancer types were integrated.

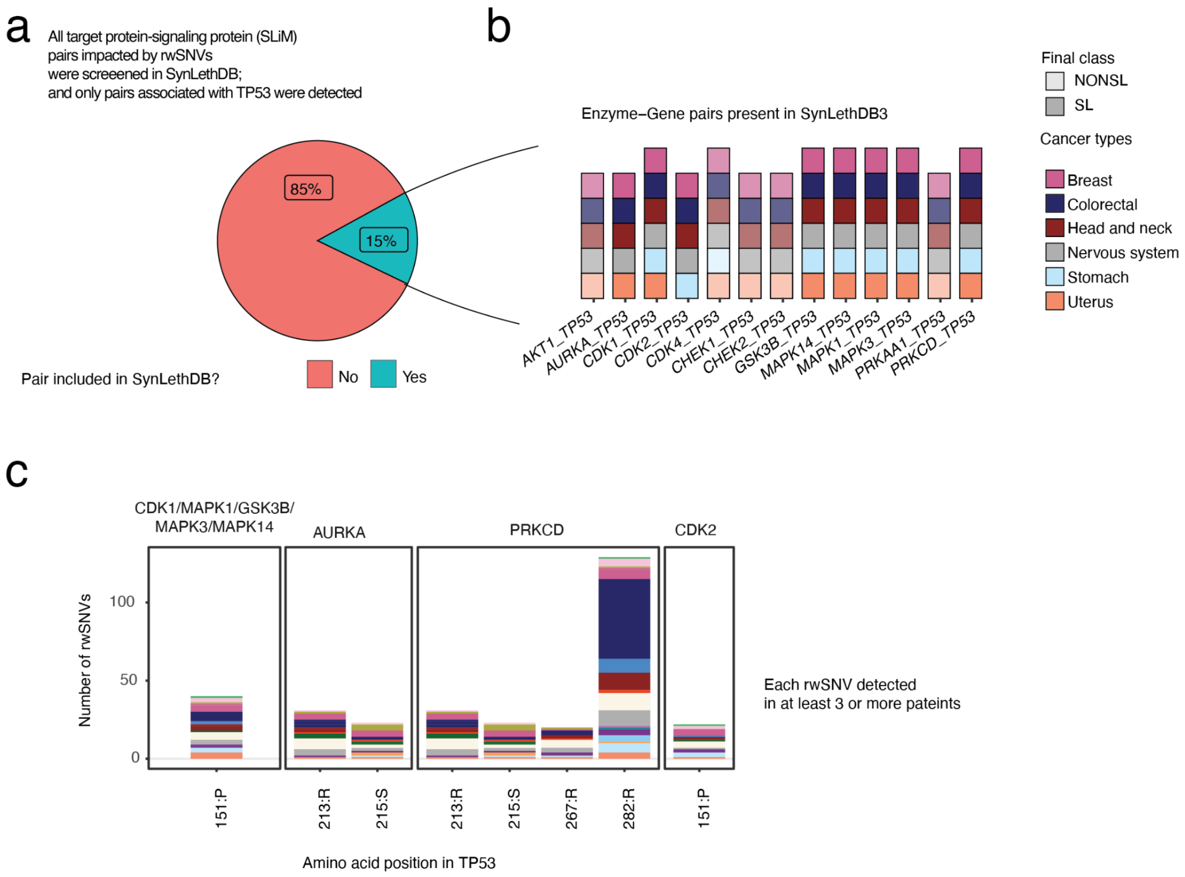


**Figure S7. Synthetic lethality of rwSNV-impacted signalling interactions involving TP53.** (**a-b**) Assessment of rwSNV-impacted target protein-signalling protein pairs for known synthetic-lethal relationships according to the SynLethDB database. Only pairs involving TP53 as the target protein were detected in the database (barplot on the right) with 8 of 13 pairs annotated as synthetic lethal. The pie chart (a) summarizes the proportion of rwSNV-impacted pairs detected in SynLethDB, and the bar plot (b) indicates the cancer types in which the rwSNV-associated interactions of TP53 and kinases were observed. (**c**) Distribution of rwSNVs in the cancer types that alter SLiMs in TP53. Bars indicate the number of rwSNVs, colors show cancer types, and facets denote the kinases that recognize the affected SLiMs. rwSNVs in SLiMs recognized by multiple kinases were counted once to reduce redundancy.
